# Differences in Characteristics of Medicare Patients Treated by Ophthalmologists and Optometrists

**DOI:** 10.1101/2019.12.31.891606

**Authors:** Darby D. Miller, Michael W. Stewart, Joshua J. Gagne, Alan L. Wagner, Aaron Y. Lee

## Abstract

**Purpose:** To quantify differences in the age, gender, race, and clinical complexity of Medicare beneficiaries treated by ophthalmologists and optometrists in each of the United States.

**Design:** Cross-sectional study based on publicly accessible Medicare payment and utilization data from 2012 to 2017.

**Methods:** For each ophthalmic and optometric provider, demographic information of treated Medicare beneficiaries was obtained from the Medicare Provider Utilization and Payment Data from the Centers for Medicare and Medicaid Services (CMS) for the years 2012 to 2017. Clinical complexity was defined using Hierarchical Condition Category (HCC) coding.

**Results:** From 2012 to 2017, ophthalmologists in every state treated significantly older beneficiaries, with the greatest difference (4.99 years in 2014) between provider groups seen in Rhode Island. In most states, there was no gender difference among patients treated by the providers, but in 46 states ophthalmologists saw significantly more racially diverse groups of beneficiaries. HCC risk score analysis demonstrated that ophthalmologists in all 50 states saw more medically complex beneficiaries and the differences were statistically significant in 47 states throughout all six years.

**Conclusions:** Although there are regional variations in the characteristics of patients treated by ophthalmologists and optometrists, ophthalmologists throughout the United States manage older, more racially diverse, and more medically complex Medicare beneficiaries.

## Introduction

Aging of the American population, with movement of the large post-world war II “baby boomers” cohort into the Medicare-covered age group, has significantly increased the prevalence of age-related eye diseases and the need for both primary and sub-specialty ophthalmology services.^1, 2^ Advancements in imaging technology, ocular pharmacotherapy, and ophthalmic surgery have broadened the spectrum of treatable disease, further increasing the total amount of eye care delivered.^2, 3^ Together, these factors suggest that a shortage of eye care providers may develop during the next 20 years. Several studies over the past 20 years have attempted to predict the future need for eye care providers, but their results have differed according to the relative proportions of provider groups used by each model.^4^

Primary care and emergency department physicians treat some patients with acute ocular conditions, but most eye care in the United States is provided by ophthalmologists and optometrists in an outpatient setting. Diagnostic procedures and treatments are performed by both professions, but because of differences in education, training, and licensure, scopes of practice differ between the two specialties both within states and between states.^5^

Not surprisingly, the geographic distribution of ophthalmologists and optometrists is not uniform within states, which has led some observers to argue that regional access problems exist for the provision of some services and the performance of some procedures.^5, 6^ Over the last several years, some states have expanded optometric scope of practice, ostensibly to improve patients’ access to eye care. Newly granted privileges include laser photocoagulation and photoablative procedures, intraocular injections, and eyelid surgeries.^5^ Data showing that these expanded privileges have improved patients’ access to care is lacking and some recent studies have concluded that access to care has not changed in states such as Oklahoma that have allowed optometrists to perform laser procedures.^5, 6, 7, 8^ Additionally, data showing that expanded privileges have enabled patients with more advanced eye diseases and higher risks of vision loss to increasingly obtain care from optometrists is also lacking. To better guide those state legislatures that are considering expanding privileges of optometrists and other non-ophthalmologists, it is critical to understand the characteristics of patients who seek care from ophthalmologists and optometrists.

To our knowledge, no published studies have evaluated the differences among Medicare beneficiaries treated by ophthalmologists and optometrists. The goal of this paper is to compare the ages, gender, race, and clinical complexity of Medicare beneficiaries seen by ophthalmologists and optometrists in all 50 states between the years 2012 and 2017.

## Methods

### Background

This cross-sectional study evaluated the 2012 to 2017 Medicare Provider Utilization and Payment Data released by the Centers for Medicare and Medicaid Services (CMS) (https://data.cms.gov) (https://www.cms.gov/Research-Statistics-Data-and-Systems/Statistics-Trends-and-Reports/Medicare-Provider-Charge-Data/Physician-and-Other-Supplier2012.html). Institutional Review Board approval was not required since the data is publicly available online and all patient data had already been de-identified.

### Identification of Ophthalmologists, Optometrists, and Patients

Ophthalmologists and optometrists were selected from the database and grouped together by provider specialty and by the National Provider Identifier (NPI). For each eye care provider and for each year in the database, we extracted name, practice address, state, and total number of unique Medicare beneficiaries. We also collected information about the average age of the beneficiaries, the number of male and female beneficiaries, the number of Caucasian beneficiaries, and the average Risk Adjustment and Hierarchical Condition Category (HCC) coding score as a measure of clinical complexity. The HCC score, which was developed by CMS^9, 10, 11^ and implemented in 2003, assigns a risk score to individuals based upon the their health conditions and demographic details. An individual’s health conditions are based on International Classification of Diseases codes submitted by providers to CMS as part of Medicare insurance claims.

Gender and racial percentages were calculated if the summed total was greater than 80% of the total reported unique Medicare beneficiaries. Otherwise, they were considered unreported. National- and state-level statistics were calculated for each demographic factor.

### Statistical Analyses

Statistical analyses were performed using R (https://www.r-project.org/) software. For each state and year, data associated with the ophthalmologists and optometrists were grouped separately to compare the average age, gender, race, and HCC risk scores for the patient cohort seen by each type of provider. Mann-Whitney U and Fisher exact tests were performed to assess differences between groups. A significance level of P < 0.05 was held to be statistically significant.

## Results

### National Data

Data for patients seen by 35,977 unique optometrists and 20,487 unique ophthalmologists from 2012 to 2017 were included in the analysis (Table 1).

**Table 1:** Demographic characteristics of Medicare beneficiaries seen by optometrists (Optom) and ophthalmologists (Ophth) between 2012 and 2017 in the United States.

|  | 2012 |  | 2013 |  | 2014 |  | 2015 |  | 2016 |  | 2017 |  | Total |  |
| --- | --- | --- | --- | --- | --- | --- | --- | --- | --- | --- | --- | --- | --- | --- |
|  | Optom. | Ophth. | Optom. | Ophth. | Optom. | Ophth. | Optom. | Ophth. | Optom. | Ophth. | Optom. | Ophth. | Optom. | Ophth. |
| **Providers (n)** | 27123 | 17402 | 27900 | 17550 | 28469 | 17665 | 28834 | 17698 | 29227 | 17788 | 29790 | 17817 | 35977 | 20487 |
| **Avg Num of Beneficiaries (n)** | 197.82 | 457.20 | 198.51 | 456.80 | 223.73 | 733.75 | 227.03 | 731.99 | 232.08 | 734.05 | 234.27 | 722.30 | 219.47 | 659.19 |
| **Avg age (years)** | 72.38 | 74.66 | 72.26 | 74.59 | 72.22 | 74.56 | 72.27 | 74.52 | 72.29 | 74.51 | 72.40 | 74.51 | 72.30 | 74.56 |
| **Avg female (%)** | 60.09 | 60.57 | 59.96 | 60.45 | 59.83 | 60.20 | 59.82 | 60.13 | 59.80 | 60.07 | 59.76 | 59.96 | 59.96 | 60.23 |
| **Avg Caucasian (%)** | 87.78 | 82.00 | 87.65 | 81.63 | 87.77 | 81.20 | 88.17 | 80.76 | 88.25 | 80.37 | 88.35 | 80.09 | 88.02 | 81.00 |
| **Age HCC Risk score** | 1.08 | 1.25 | 1.06 | 1.22 | 1.03 | 1.20 | 1.09 | 1.26 | 1.09 | 1.28 | 1.09 | 1.29 | 1.07 | 1.25 |

Each ophthalmologist saw an average of 439.7 more Medicare beneficiaries each year compared to each optometrist. Nationally, ophthalmologists cared for older beneficiaries than did optometrists (74.45 versus 72.30 years; P < 0.0001). The majority of the beneficiaries were female and the proportion female was similar between optometrists (59.96%) and ophthalmologists (60.23%). Ophthalmologists treated a more racial diverse patient population than did optometrists and cared for a significantly higher percent of non-Caucasian patients (total average Caucasian: 88.02% for optometrists and 80.10% for ophthalmologists, P < 0.0001). Patients seen by optometrists were significantly healthier with lower average HCC risk scores than those seen by ophthalmologists 1.07 vs. 1.25 (P < 0.0001).

### State Age Data

In all states and during all years, ophthalmologists saw significantly older patients compared to optometrists (Figure 1 and Supplementary Figure 1). The youngest average patient population (70.01 years, 95% CI: 69.53 to 70.49 years) was seen by optometrists in the state of Louisiana in 2015 and the oldest average patient population (77.10 years, 95% CI: 76.41 to 77.79 years) was seen by ophthalmologists in North Dakota in 2015. The largest difference in ages between men and women was in Rhode Island in 2014 (female – male = 4.99 years, 95% CI: 4.08 to 5.90 years).

**Figure 1.**
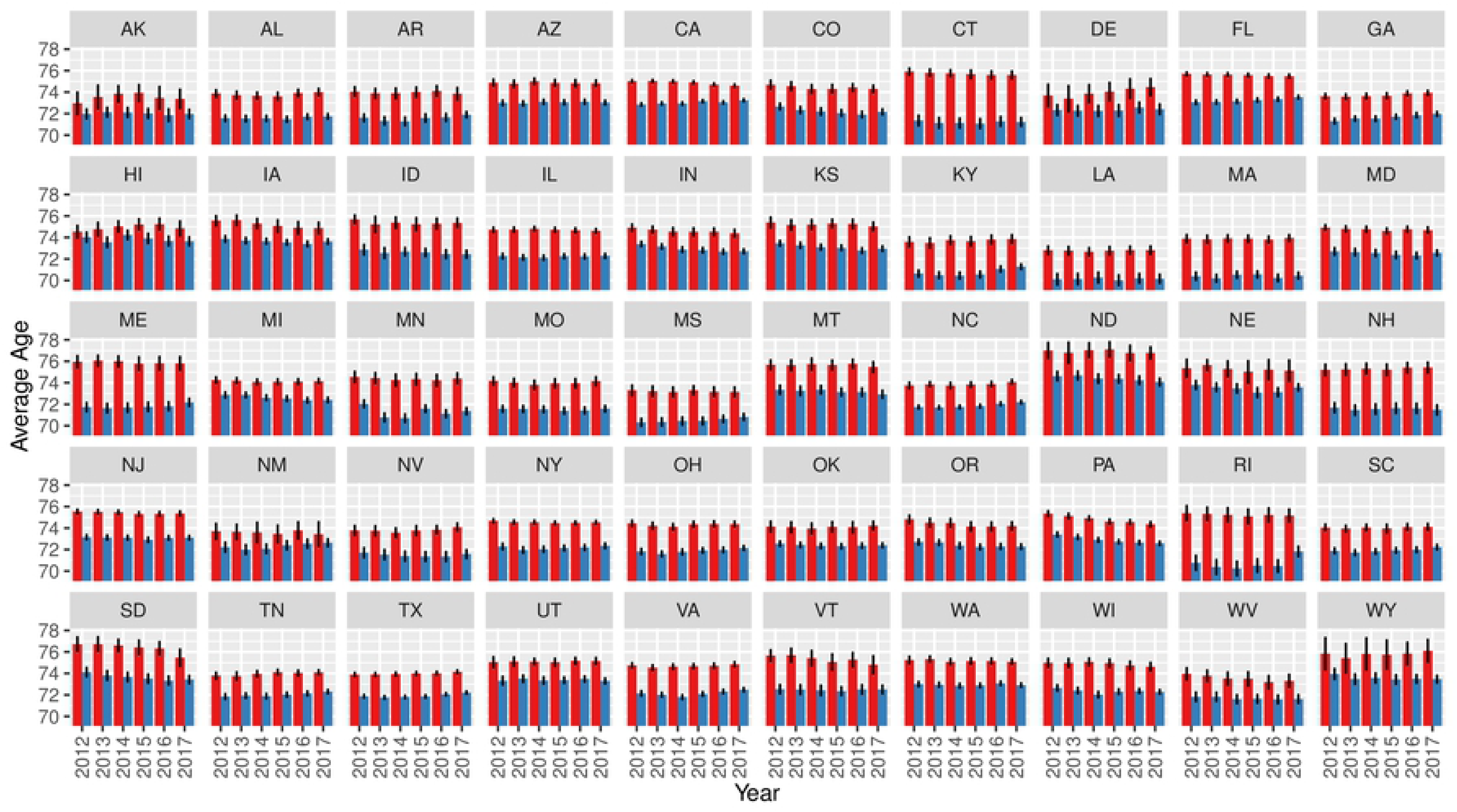
Average age of Medicare beneficiary population by state and eye provider type. Ophthalmologists are shown in red, and optometrists are shown in blue. The error bars represent 95% confidence intervals.

### State Gender Data

In 41 of the 50 states, there was no significant difference in the genders of patients seen by eye care providers from 2012 to 2017 (Figure 2 and Supplementary Figure 2). In Minnesota, New Jersey, New York, Wisconsin, Connecticut, and Pennsylvania, ophthalmologists saw more females than did optometrists during all six years. In Alabama, Oklahoma, and Mississippi, this trend was reversed, as optometrists saw more females than did ophthalmologists.

**Figure 2.**
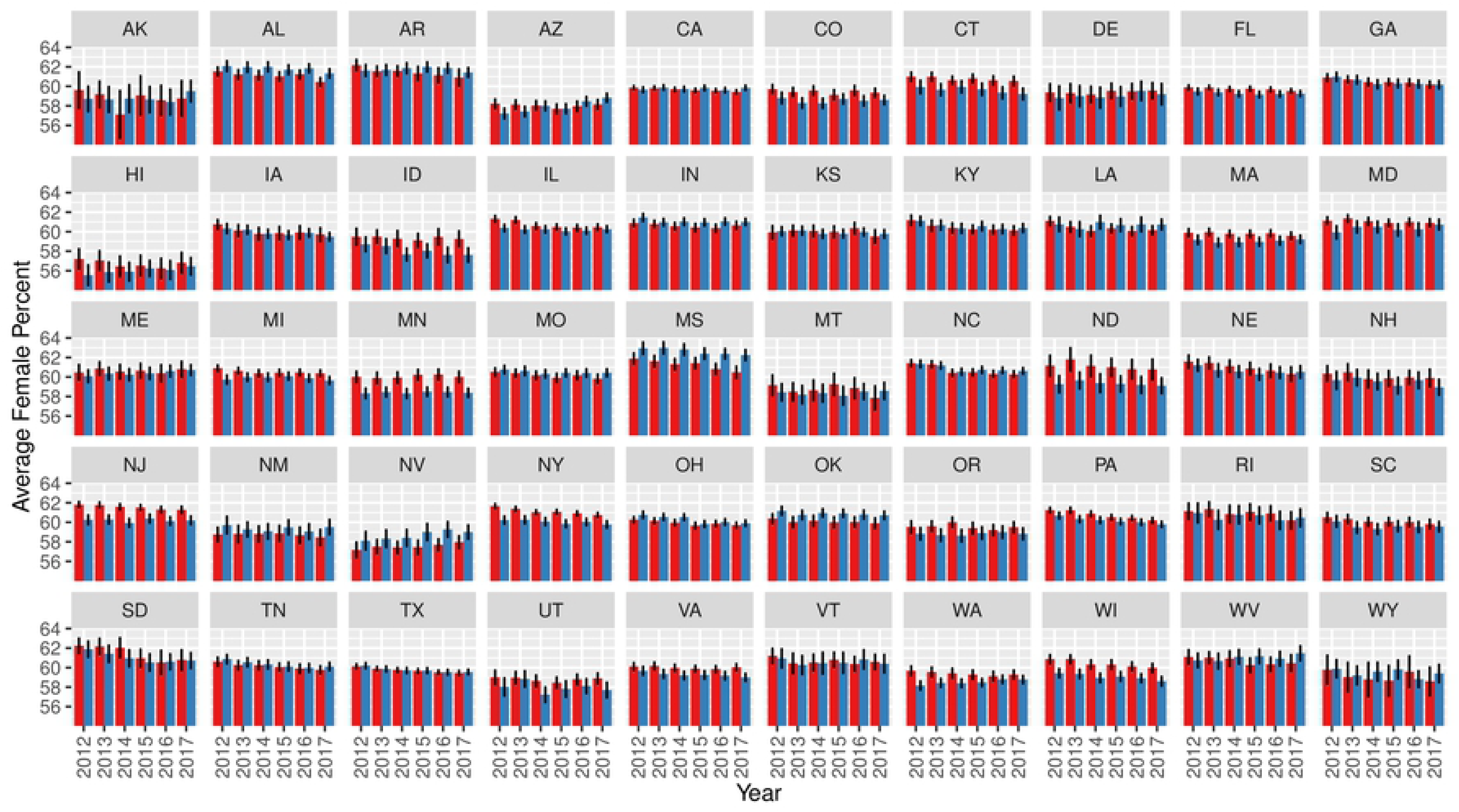
Average percentage of female Medicare beneficiary population by state and eye provider type. Ophthalmologists are shown in red, and optometrists are shown in blue. The error bars represent 95% confidence intervals.

### State Racial Data

In 42 states during each of the six years, there was a significant difference in the racial composition of the beneficiaries seen by ophthalmologists and optometrists (Figure 3 and Supplementary Figure 3). In every state except for Arizona, Montana, North Dakota, and South Dakota, ophthalmologists saw more non-Caucasian patients than did optometrists. The states without a significant difference were Hawaii, New Mexico, Louisiana, Mississippi, Alaska, Delaware, Rhode Island, and Connecticut.

**Figure 3.**
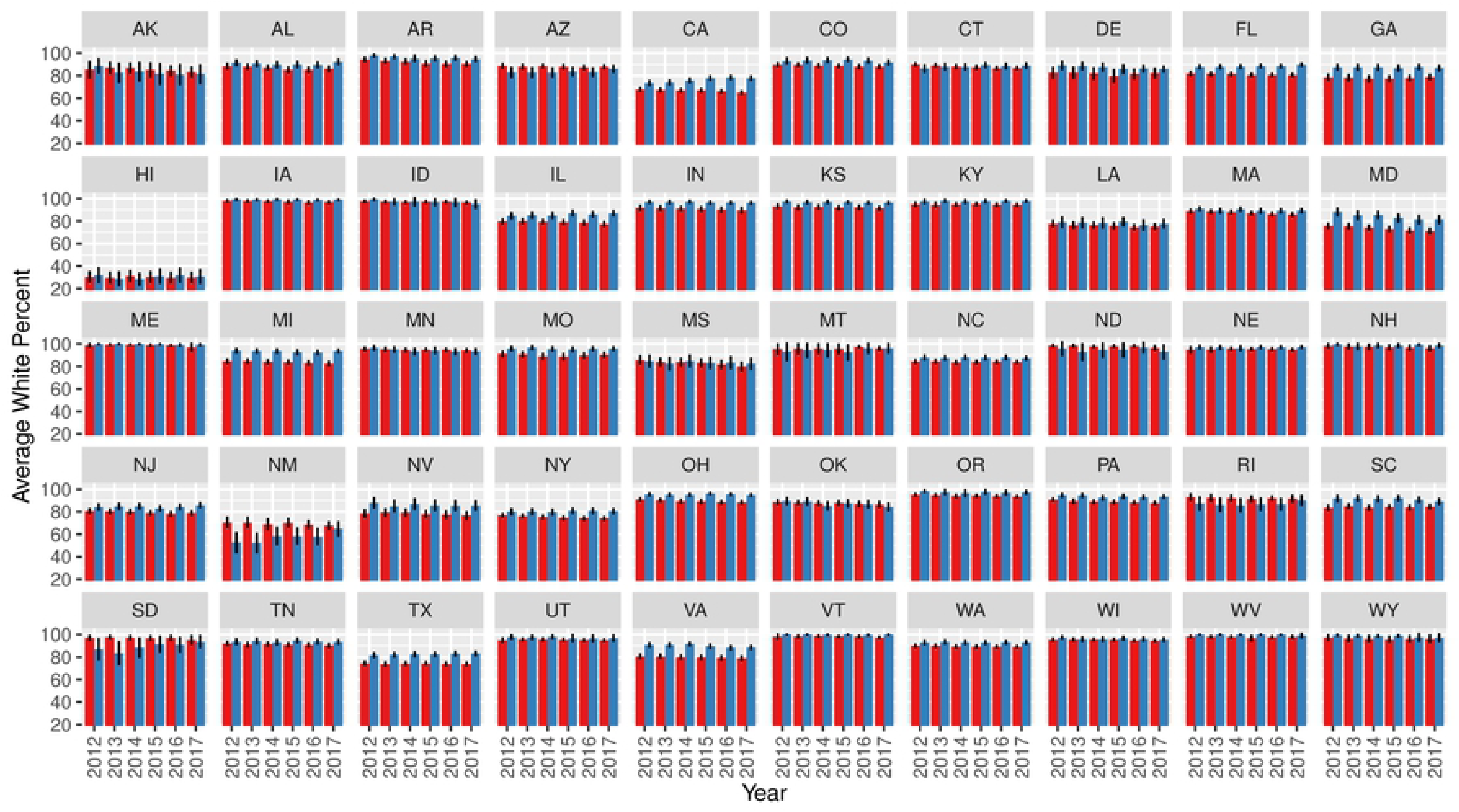
Average percentage of Caucasian Medicare beneficiary population by state and eye provider type. Ophthalmologists are shown in red, and optometrists are shown in blue. The error bars represent 95% confidence intervals.

### State HCC Data

During every year in every state, ophthalmologists treated more medically complex patients compared to optometrists (Figure 4). This trend was statistically significant in every state except for Mississippi, Wyoming, and Alaska (Supplementary Figure 4). The greatest difference was in Vermont in 2017 with ophthalmologists seeing 32.28% more complex patients (95% CI: 22.12% to 42.44%). Scope of practice laws changed in Louisiana in 2014, allowing optometrists to perform certain types of eye and eyelid surgery. When HCC scores for optometrists before and after 2014 were compared, there was no statistically significant difference (P = 0.22). There was, however, a statistically significant increase in HCC scores for ophthalmologists in Louisiana when comparing after 2014 to before 2014 (P < 0.05) indicating an increase in clinical complexity of patients being seen.

**Figure 4.**
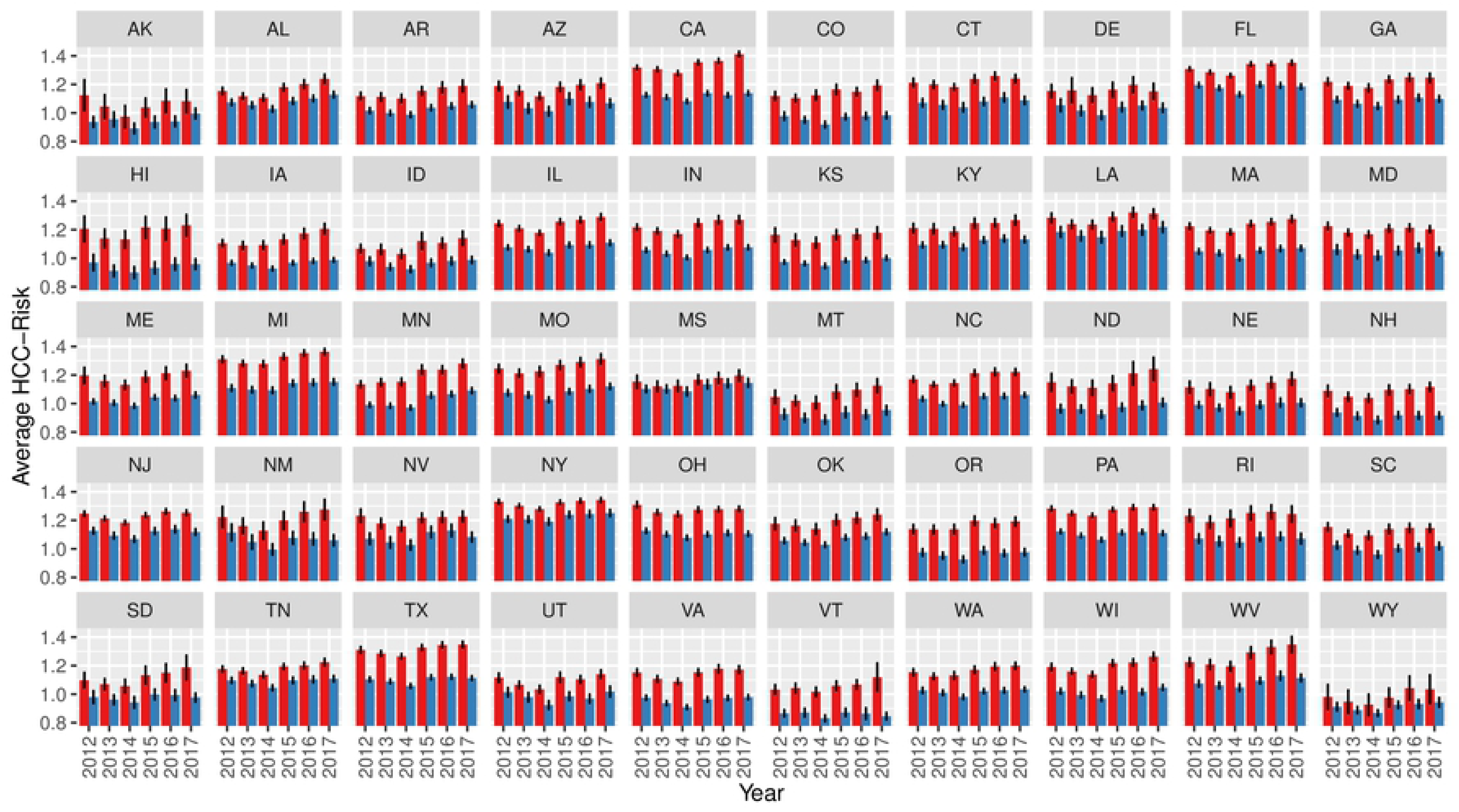
Average Hierarchical Condition Category (HCC) risk score by state and eye provider type. Ophthalmologists are shown in red, and optometrists are shown in blue. The error bars represent 95% confidence intervals.

## Discussion

To the best of our knowledge, this is the first study to compare patients seen by ophthalmologists and optometrists according to their ages, gender, race, and clinical complexity. According to Medicare payment and utilization data from 2012 to 2017, it is evident that ophthalmologists see older, sicker, and more racially diverse patients than do optometrists.

Hierarchical Condition Category Coding was introduced by CMS as a risk-adjustment approach to estimate future health care costs for patients.^6^ HCC coding relies on ICD coding to assign risk scores to patients. The ICD codes and additional risk factors for future health care costs, such as gender, age, living situation and Medicaid eligibility, are used to calculate a Risk Adjustment Factor (RAF) that is assigned to each HCC category. As the healthcare landscape has increasingly shifted to value-based payment models, HCC coding has become more prevalent. Insurance companies often use the RAF to predict future health care costs. A patient with few serious health care conditions would have a low RAF but a patient with several chronic conditions would be expected to have greater health care utilization and incur higher health care costs. Patients who are healthier than average will have a HCC score less than 1.00, whereas those that are less healthy than average have HCC scores greater than 1.00. Overall, HCC coding gives a single view of each patient by representing complexity.

The reasons that ophthalmologists see patients with higher HCC scores are not fully known but several factors can be considered. The incidences of the most common vision threatening conditions in the United States (cataracts, age-related macular degeneration, glaucoma, diabetic retinopathy) increase with patient age; therefore, it would be expected that elderly patients have higher HCC risk scores. It may be that many elderly patients recognize that ophthalmologists, with their medical school education, and medical and surgical residency training, are better equipped than optometrists to prescribe pharmacotherapies and perform surgeries for complex conditions.

Many patients with age-related vision threatening diseases may have presented initially to optometrists or general physicians but required referral to ophthalmologists for more advanced evaluation and treatments. Funneling of complex patients to ophthalmologists would be expected to increase the HCC scores among ophthalmologic practices.

Optometric practices are often associated with retail optical chains, some of which are located within shopping malls. These may attract younger, more mobile patients than ophthalmology practices, many of which are located in multi-specialty medical centers or on hospital campuses. Optometrists often market their practices based on expertise in contact lenses and glasses whereas ophthalmologists often market practices based on surgery.

Several strengths and weakness of this study need to be mentioned. The study has the inherent weaknesses of a retrospective study, including data that was collected after the encounters. As with any secondary database analysis, the validity of the results depends on the accuracy of the underlying data. Data for all variables were not fully reported for each state in each year. Moreover, ICD codes from Medicare claims data may not capture with perfect sensitivity the comorbid conditions that patients have. We examined patient age, gender, race, and HCC risk score averages by state, but not at more granular county, city, or individual practitioner levels. Race data are only broken down into Caucasian and non-Caucasian. The strength of the study lies in the large numbers of practitioners and patients identified from the large cohort of United States Medicare beneficiaries.

The data from this study show that ophthalmologists and optometrists see patient populations that differ with respect to age and disease severity. On average, ophthalmologists have more experience dealing with elderly patients with more severe medical conditions. Health policy makers should understand that the two professions cannot be viewed interchangeably as some would suggest and that decisions to expand access to patient care by legislatively removing scope of practice barriers should be considered carefully. The data shows that moving complicated patients from ophthalmologists to optometrists would shift their care to less experienced providers. Further, any studies that compare outcomes of care provided by ophthalmologists to optometrists will need to carefully account for the differences in patient characteristics that we observed.

---

**Supplementary Figure 1:**
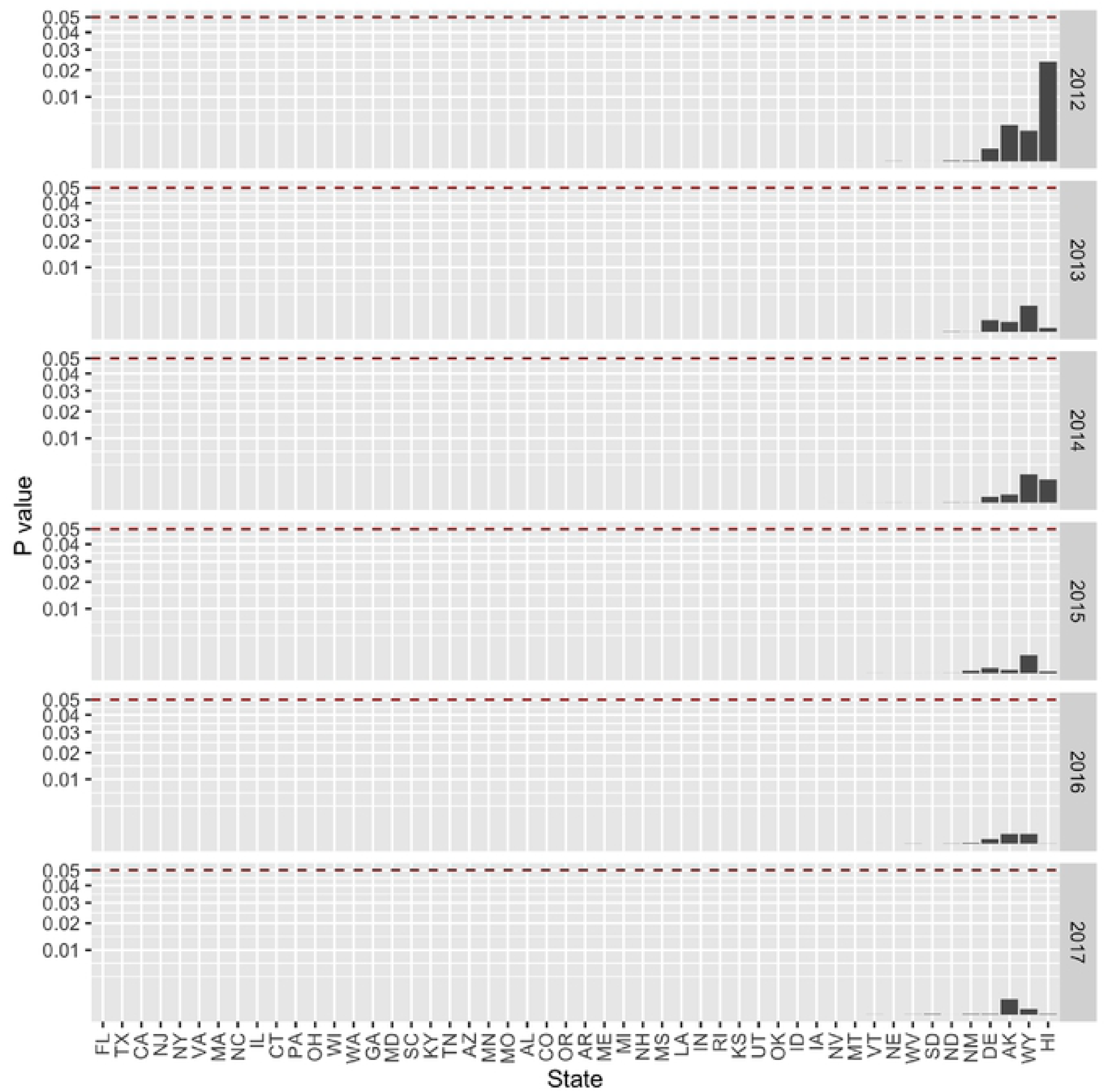
Statistical significance test for average age comparing differences between ophthalmologists and optometrists for each state and year. The red dashed line represents the threshold for statistical significance. The lowest possible value is 2.2 x 10^-16^ and most of the P values are too small to be seen on this scale.

**Supplementary Figure 2:**
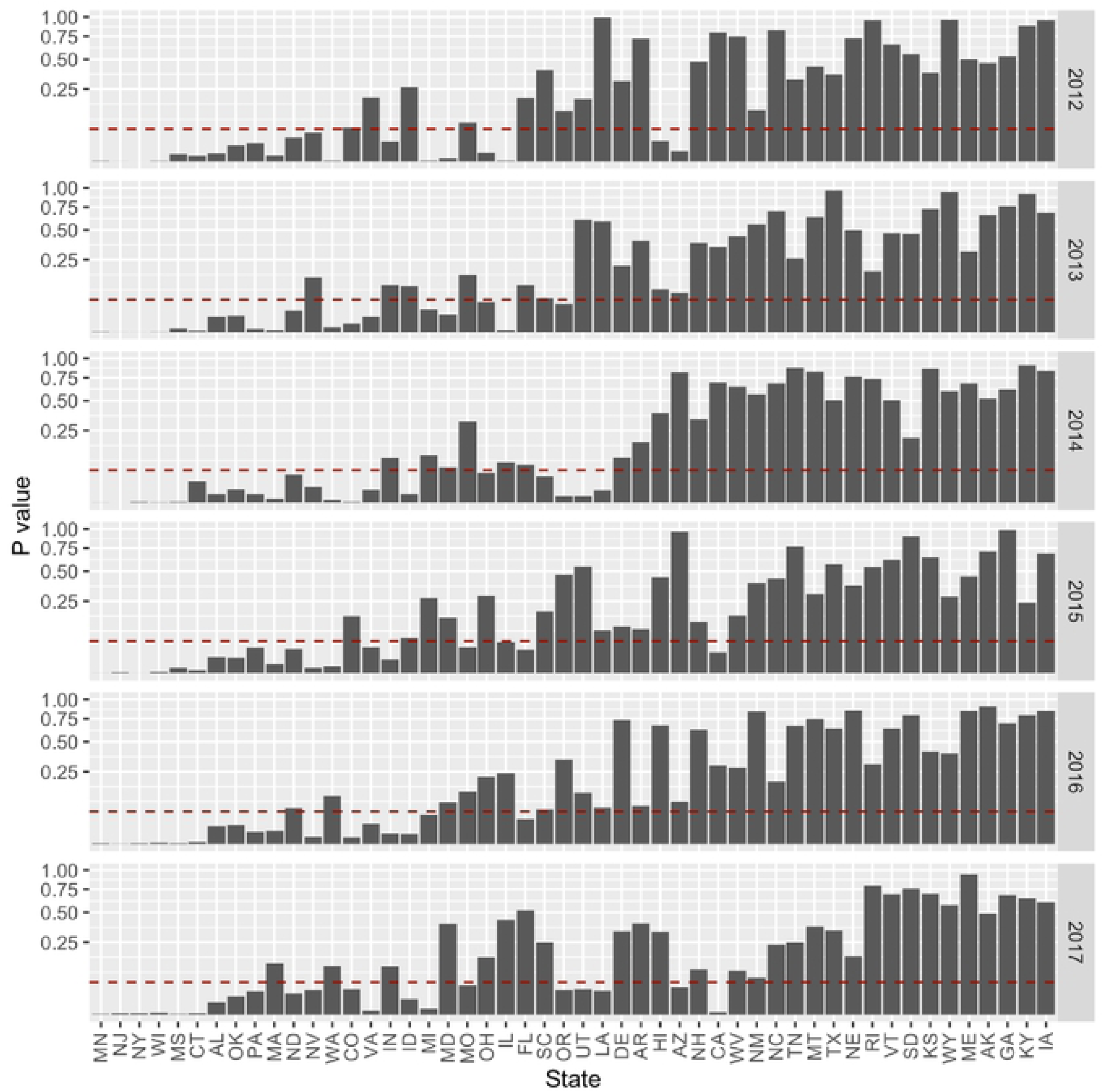
Statistical significance for percent of female beneficiaries comparing differences between ophthalmologists and optometrists for each state and year. The red dashed line represents the threshold for statistical significance. The lowest possible value is 2.2 x 10^-16^.

**Supplementary Figure 3:**
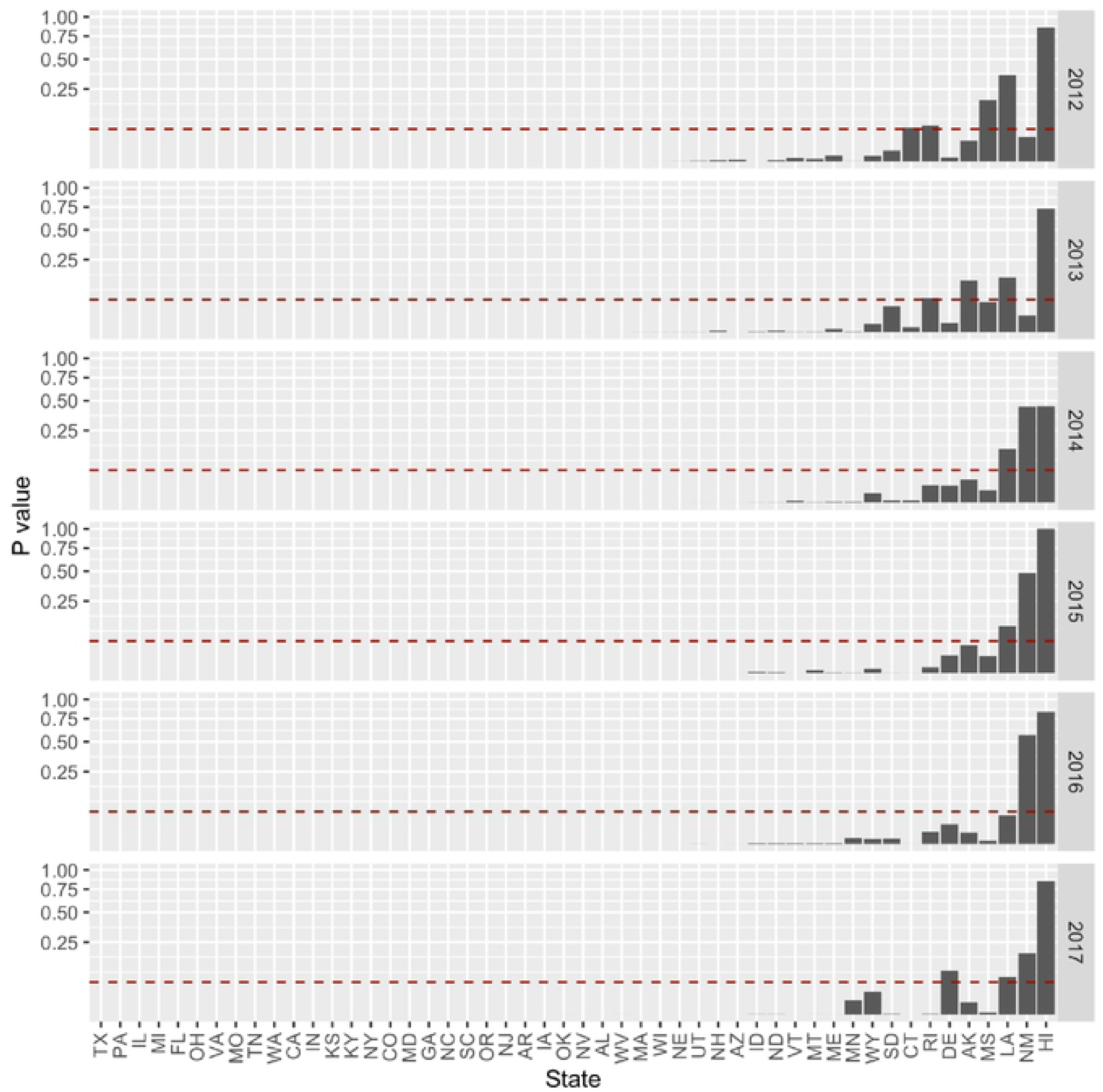
Statistical significance for percent of Caucasian beneficiaries comparing differences between ophthalmologists and optometrists for each state and year. The red dashed line represents the threshold for statistical significance. The lowest possible value is 2.2 x 10^-16^ and most of the P values are too small to be seen on this scale.

**Supplementary Figure 4:**
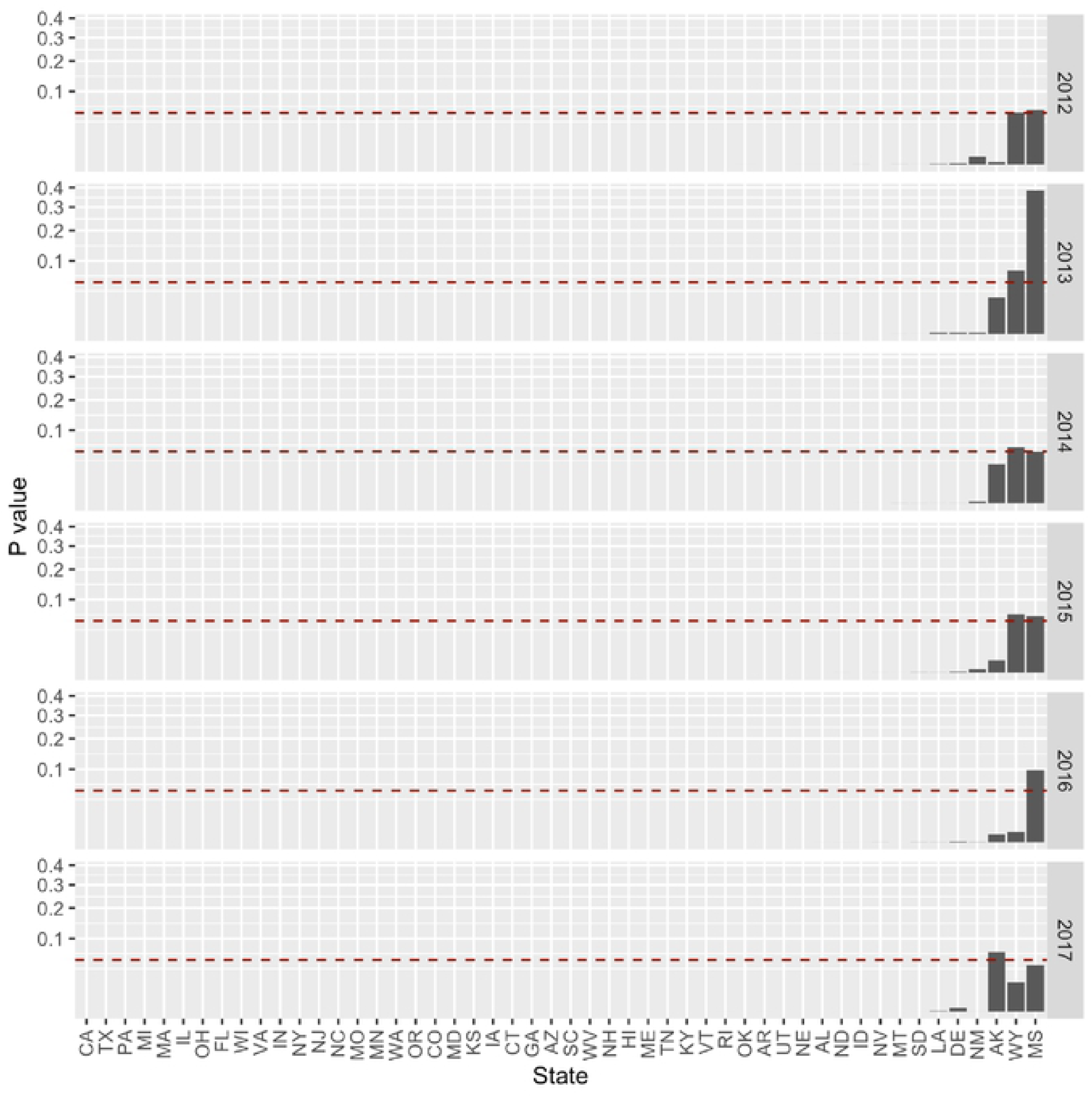
Statistical significance for Hierarchical Condition Category risk score comparing differences between ophthalmologists and optometrists for each state and year. The red dashed line represents the threshold for statistical significance. The lowest possible value is 2.2 x 10^-16^ and most of the P values are too small to be seen on this scale.

